# Biosynthesis and apoplast accumulation of the apocarotenoid pigment azafrin in parasitizing roots of *Escobedia grandiflora*

**DOI:** 10.1101/2022.02.07.479377

**Authors:** Edison Cardona-Medina, Marisa Santos, Rubens Nodari, Damaso Hornero-Méndez, Arnau Peris, Darren C. J. Wong, José Tomás Matus, Manuel Rodríguez-Concepción

## Abstract

- The herbaceous hemiparasite *Escobedia grandiflora* (Orobanchaceae) is used in traditional medicine in the Andean region. Their roots accumulate an orange pigment with a significant relevance as a cooking dye that exhibits antioxidant and cardioprotective properties.
- The present work combined metabolic and cytological analyses with *de novo* transcriptome assembly, gene expression studies, and phylogenetic analyses to confirm the chemical identity of the pigment and investigate its biosynthesis and function in *Escobedia* roots.
- The pigment was conclusively shown to be azafrin, an apocarotenoid likely derived from the cleavage of β-carotene. Candidate genes for the production of azafrin in *Escobedia* roots are proposed based on RNA-seq supported by RT-qPCR and phylogeny reconstruction analyses. In particular, our data suggest that azafrin production relies a carotenoid cleavage dioxygenase (CCD) different from CCD7 and similar to CCD4 enzymes. We also show that azafrin is delivered to the root apoplast and that it accumulates in the area where the *Escobedia* haustorium contacts the host’s root, suggesting a role of azafrin in the parasitization process.
- Altogether, our work represents an unprecedented step forward in our understanding of the *Escobedia* parasitization system, but it also provides vital information towards the eventual domestication of this valuable medicinal plant.

## Introduction

*Escobedia grandiflora* (L. f.) Kuntze (Orobanchaceae) (hereafter referred to as *Escobedia*) is a perennial hemiparasitic plant native to Central and South America, where it associates with high diversity in plant communities of dry and wetland non-forested ecosystems (Burguer & Barringer, 2000; Cardona & Muriel, 2015; Cardona-Medina *et al*., 2019; Cardona-Medina *et al*., 2021). Wild populations of this herbaceous plant have supported several traditional uses in the Andean region. The abundant orange pigment in its root is used as a cooking dye and local medicine for hepatitis, jaundice, hyperlipidaemia, and obesity (Silva *et al*., 2010; Muriel *et al*., 2015). Such water-soluble pigment was tentatively proposed to be azafrin, a C27 apocarotenoid (Kuhn, 1935; Eschenmoser & Eugster, 1975). Azafrin, which exhibits antioxidant and cardioprotective properties (Yang *et al*., 2018), is found at high levels in particular organs of other parasitic plants, including the roots of *Centranthera grandiflora* and the rhizomes of *Alectra parasitica*, but also in non-parasitic medicinal plants such as *Caralluma umbellata* (Agrawal *et al*., 2014; Evanjaline & Vr, 2018; Verma *et al*., 2019; Zhang *et al*., 2019).

Apocarotenoids are cleavage products of carotenoids. Some of them are biologically active molecules with roles as regulators of plant development and environmental interactions, including abscisic acid (ABA), strigolactones (SL), and others yet to be fully characterized (Walter *et al*., 2010; Hou *et al*., 2016; Felemban *et al*., 2019; Moreno *et al*., 2021). Plant apocarotenoids often modulate the interaction with herbivores, arbuscular mycorrhizal (AM) fungi, and parasitic plants (Moreno *et al*., 2021; Wang *et al*., 2021). A well-known case is SL, which stimulate the germination of some parasitic seeds from Orobanchaceae, contribute to establishing AM symbiosis, and modulate other rhizospheric communication with symbionts and parasites (Torres-Vera *et al*., 2016; Mutuku *et al*., 2021). Other apocarotenoids with roles in rhizospheric interactions include blumenols, mycorradicins, zaxinone and anchorene (Moreno *et al*., 2021; Wang *et al*., 2021).

Biosynthesis of plant carotenoids takes place in plastids, and it begins with the formation of C_40_ 15-*cis*-phytoene through condensation of two molecules of C_20_ geranylgeranyl diphosphate (GGPP) by phytoene synthase (PSY), the first and main rate-determining enzyme of the pathway (Rodriguez-Concepcion *et al*., 2018; Moreno *et al*., 2021). PSY is usually encoded by small gene families in plants (Stauder *et al*., 2018). Phytoene desaturation and isomerization produce lycopene, the red carotenoid responsible for the colour of ripe tomatoes. The formation of β rings in the two ends of the lycopene molecule generates β-carotene, the main pro-vitamin A carotenoid and the proposed precursor of azafrin (Rodriguez-Concepcion *et al*., 2018; Zhang *et al*., 2019). Cleavage of the C_40_ skeleton of carotenoids to produce apocarotenoids can either take place non-enzymatically or be catalysed by carotenoid cleavage dioxygenase (CCD) enzymes (Felemban *et al*., 2019; Moreno *et al*., 2021). In the model plant *Arabidopsis thaliana*, CCD enzymes of the 9-*cis*-epoxycarotenoids dioxygenase (NCED) type are involved in the biosynthesis of ABA whereas CCD7 and CCD8 participate in the first steps of SL production. Other CCD enzymes such as plastid-localized CCD4 and cytosolic CCD1 are involved in the production of other apocarotenoids, including volatiles and growth regulators (Auldridge *et al*., 2006; Hou *et al*., 2016; Felemban *et al*., 2019; Moreno *et al*., 2021). The biosynthetic pathway of azafrin was proposed to begin with the isomerization of β-carotene to 9-*cis*-β-carotene by the enzyme DWARF27 (D27), followed by cleavage by CCD7 to produce 10’-apo-β-carotenal, a common precursor of SL (Bruno & Al-Babili, 2016; Zhang *et al*., 2019). Then, an unknown aldehyde dehydrogenase would transform aldehyde into carboxylic acid, and a cytochrome P450 monooxygenase would catalyse the oxidation reactions to produce azafrin (Zhang *et al*., 2019).

Here we use modern mass-spectrometry technologies to ascertain the identity of the *Escobedia* root pigment as azafrin, propose candidate genes involved in its production, demonstrate its accumulation in the apoplastic space of root cells, and provide insights on its possible function in parasitic plant-host interactions.

## Materials and Methods

### Plant material

Roots and dried fruits of *Escobedia grandiflora* (L. f.) Kuntze were collected from nine mature individuals (i.e., in the flowering period) in a wild population located in the Lagoinha do Leste Municipal Park (27°46’53.0”S, 48°29’16.1”W, 190 m.a.s.l.), municipality of Florianópolis, Brazil. After collection, the roots were frozen at −80°C, lyophilized, and then pulverized with a TissueLyser (Qiagen) to obtain a fine powder for further chromatography and RNA extraction. Seeds were collected from dried fruits, imbibed for five days in distilled water and sown as described by Cardona-Medina *et al*. (2019). Twenty imbibed seeds were sown in 2-liter pots without a host plant in greenhouse conditions with a long day photoperiod. After nine months, roots were collected, frozen and processed as described above for wild (host-attached) samples. Furthermore, twenty imbibed seeds were sown in 5-liter pots with *Pennisetum purpureum* Schumach (Poaceae) as a host plant (Cardona & Muriel, 2015; Cardona-Medina *et al*., 2019) and grown for nine months to observe different root and haustoria development phases, as reported by Cardona-Medina *et al*. (2019).

### Azafrin identification and quantification analysis

Azafrin identification was based on HPLC-DAD-MS(APCI+) analysis carried out as described (Supporting Information Methods S1). For azafrin quantification, lyophilized roots from *Escobedia* plants were used for pigment extraction, chromatographic separation, and detection at 450 nm as described (Supporting Information Methods S1). Three biological replicates were performed in every experiments.

### RNA-seq

Total RNA was extracted from a pool of *Escobedia* roots grown with their hosts in a wild population. Samples representing different stages of the parasitizing process (i.e. root development) were extracted using the Maxwell® RSC Plant RNA kit (Promega) with the Maxwell® RSC Instrument (Promega), according to the manufacturer’s instructions. RNA samples were pooled in equivalent amounts and library construction was performed with Illumina TruSeq Stranded mRNA kit, according to the manufacturer’s instructions. A single library was sequenced at a depth of 30 million reads. Raw paired-end (2×150 bp) reads were processed as follows. Removal of adaptor sequences, sliding-window trimming, length, and quality filtering of the raw paired-end reads were performed with *fastp* v0.20.0 (Chen *et al*., 2018) using *-w* 16 −5 −3 *-r -W* 4 *- M* 20 *-l* 40. All other settings were left at default. *De novo* transcriptome assembly of surviving reads was performed with *Trinity* v2.11.0 (Haas *et al*., 2013) using default settings. Transcriptome analysis and functional annotation was performed as described (Supporting Information Methods S2).

### Phylogenetic analysis

Phylogenetic reconstructions were performed with MrBayes (Huelsenbeck & Ronquist, 2001). The resulting topology based on a phylogram was constructed through the standard stepwise pipeline. Firstly, the candidate protein sequences were aligned with MUSCLE (Edgar, 2004) using default parameters and the multiple sequence alignment outcome was exported in nexus format (Maddison *et al*., 1997). MEGAX (Kumar *et al*., 2018) was used to identify the best substitution protein model, i.e. with the lowest Bayesian information criterion (BIC) score. Each sequence was considered as ‘partially-deleted’ in which a candidate is discarded if it presents higher percentage of ambiguous sites than the threshold specified in the Site Coverage Cutoff parameter in MEGAX options (similar to the complete deletion option but with a threshold set to 100%, i.e. absence of ambiguous sites). Once the multiple sequence alignment was performed, the best substitution model was identified and the distribution of rates among sites was selected. In our case, the best model was Jones-Taylor-Thornton (JTT) with rates among sites following a gamma distribution. The phylogenetic analysis was then conducted by bayesian inference, using the Markov Chain Monte Carlo algorithm (MCMC) available in MrBayes. The different clusters’ reliability was calculated with the posterior probabilities (Gelman *et al*., 1995), being the values higher than 0.70 acceptable and the values higher than 0.90 highly supported.

### RT-qPCR

RNA extracted from *Escobedia* roots was used for real-time quantitative PCR (RT-qPCR) as described (Supporting Information Method S3). RT-qPCR data were normalized with the *Escobedia* actin gene DN8798_c0_g1_i1. Primers are listed in Supporting Information Table S1. For statistical analyses of the results, a one-way ANOVA was performed in which the host conditions were the explanatory variable, and azafrin and relative transcript levels of candidate genes were the response variable. Response variables were square-root transformed. ANOVA validation was based on the Shapiro-Wilk test and the analysis of plotting residuals. Significant differences (p≤0.05) were tested using Tukey’s test. The data were analyzed in R environment version 4.0.3 (R Core Team, 2021).

### Structural and anatomical analyses

Anatomical analyses were performed as described (Supporting Information Methods S4). Haustoria were fixed in a solution with 2.5 % glutaraldehyde in a 0.1 M sodium phosphate buffer and dehydrated through an ethanolic series (Ruzin, 1999). For scanning electron microscopy (SEM) analyses, the fixed and dehydrated samples were processed as described (Supporting Information Methods S4). The hyaline body ultrastructure was visualized by transmission electron microscopy (TEM), according to Pueschel (1979) after processing the samples as described (Supporting Information Methods S4). Azafrin detection by confocal microscopy of fresh hand-cut root sections was performed as described by D’Andrea *et al*. (2014), using the 488 nm ray line of an argon laser for excitation and 500-550 nm of emission window.

## Results and Discussion

### Escobedia grandiflora roots accumulate azafrin and also produce aeginetin

While early studies in *Escobedia* species proposed that the orange pigment that accumulates in roots is azafrin (Kuhn, 1935; Karrer & Jucker, 1948; Eschenmoser & Eugster, 1975), clearcut demonstration is still missing. Consistent with those studies, the HPLC analysis of a root extract from *Escobedia* plants of a wild population revealed the presence of a major compound (>95%; peak 2) with an on-line UV-visible spectrum with maxima at 390, 421, 436 nm (Fig. 1). This spectrum agrees with a chromophore structure with seven to eight conjugated double bonds (c.d.b), in accordance with the structure proposed for azafrin with a seven c.d.b. polyene chain and a conjugated carbonyl group (Fig. 1a). However, the acid mobile phase likely affected absorption maxima and fine structure, resulting in the UV-visible spectrum being slightly different from that reported in the literature (Britton, 1991; Britton *et al*., 2004). When peak 2 was collected and purified from the diode array detector outlet, the resulting UV-visible spectrum in ethanol (388, 407, 431 nm) matched the one reported for azafrin in previous studies (Britton, 1991; Britton *et al*., 2004) (Fig. 1b). The chemical identity of this peak was confirmed by mass spectrometry using HPLC-DAD-MS(APCI+). As shown in (Fig. 1c), the mass spectrum was consistent with the formula C_27_H_38_O_4_ (MW=426.2770), with a fragmentation pattern presenting three characteristic fragments corresponding to the protonated molecule ([M+H]+, 427.27) and the loss of one and two water molecules derived from the hydroxy groups ([M+H-18]+, 409.26; [M+H-18-18]+, 391.25). The fragment derived after the neutral loss of the carboxylic group [M-46]+ at 381.26 was also detected but with low relative abundance. These results conclusively confirmed that peak 2 of the chromatogram, corresponding to the most abundant pigment by far in *Escobedia* roots (Fig. 1a), was indeed azafrin.

**Fig. 1.**
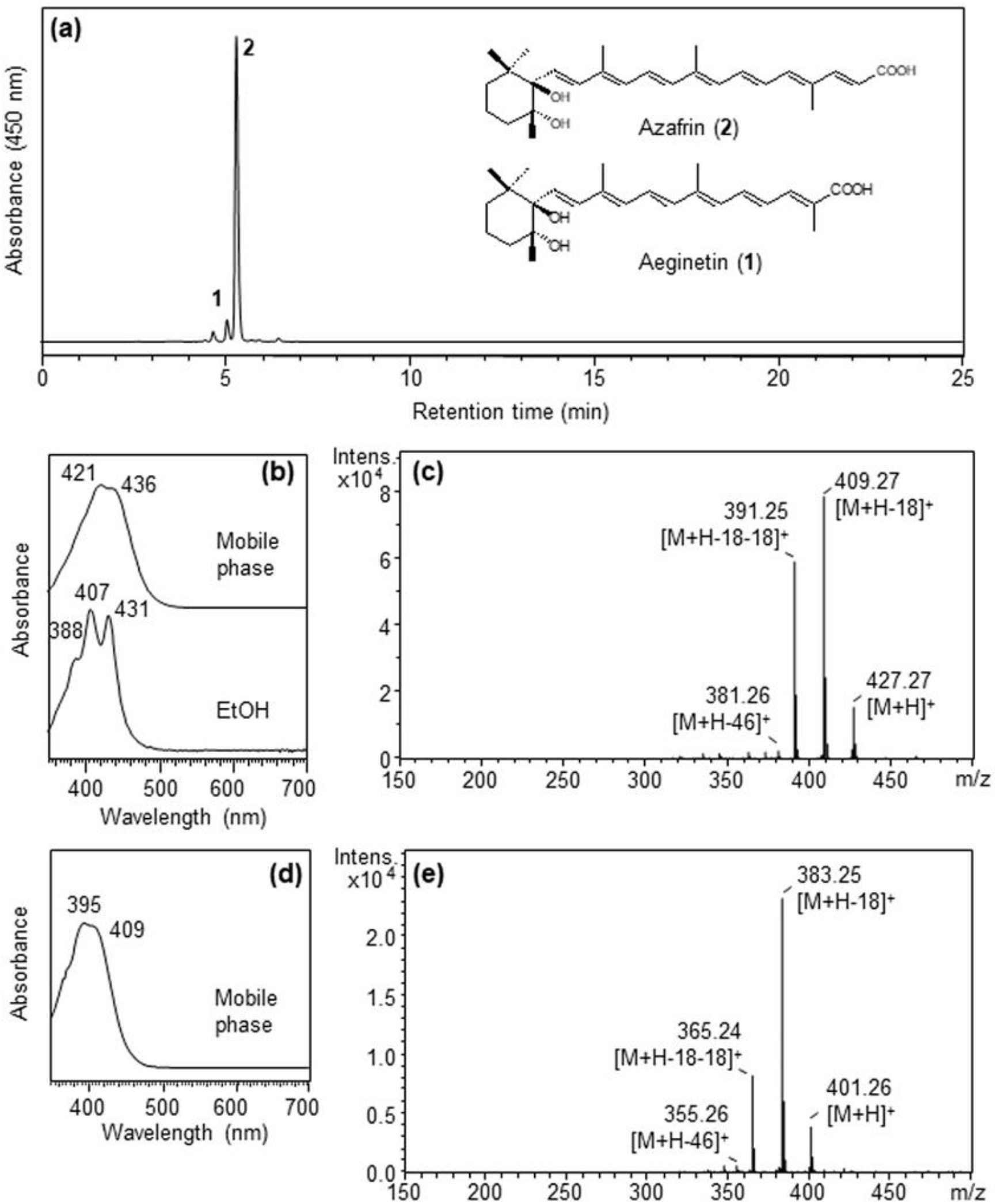
*Escobedia* roots accumulate azafrin and lower levels of aeginetin. **(a)** HPLC-DAD chromatogram of *Escobedia* root pigment extract at 450 nm; **(b)** UV/Vis spectra of the major peak (**2**; azafrin) in the mobile phase and after isolation in ethanol; **(c)** mass spectrum of peak **2**; **(d)** UV/Vis spectra of the minor peak (**1**; aeginetin) in the mobile phase; **(e)** mass spectrum of peak **1**.

Regarding the minor compound eluting before azafrin (peak 1; Fig. 1a), its UV-visible spectrum presented maxima at 395 and 409 nm (Fig. 1d), suggesting a chromophore with 6 c.d.b. (i.e., 1 c.d.b. shorter compared to azafrin). The mass spectrum of this compound showed characteristic fragments: [M+H]+ at 401.26, [M+H-18]+ at 383.25, [M+H-18-18]+ at 365.24 and [M-46]+ at 355.26 (Fig 1e). Remarkably, this mass fragmentation profile was very similar to the one observed for azafrin but fragments showed 28 u.m.a. less, which correspond to an alkene unit (-CH_2_=CH_2_-). These data corroborated that the structure has a chromophore with 1 c.d.b less than azafrin. The protonated molecule ([M+H]+ at 401.26) was consistent with a molecular structure with a formula C_25_H_36_O_4_ (MW=400.2613), which corresponds to the apocarotenoid aeginetin (Fig. 1a), isolated from the roots of *Aeginetia indica* (Eschenmoser *et al*., 1982; Britton, 1991), a holoparasitic herb of the same plant family as *Escobedia*, Orobanchaceae.

Azafrin has been identified in non-parasitic species but it is only accumulated at high levels in the roots or rhizomes of root hemiparasitic plants (Agrawal *et al*., 2014; Zhang *et al*., 2019). It is interesting to note that azafrin-overaccumulating hemiparasitic plants such as *Escobedia*, *Centranthera grandiflora* (hereafter referred to as *Centranthera*) and *Alectra parasitica* are classified in the same subclade (I) within the Buchnerae clade of Orobanchaceae family (Nickrent, 2020). Two other species belonging to this subclade I (*Melasma stricta* and *Notochilus coccineus*) were reported to have orange roots, but the identity of the pigment has not been identified yet (Safford, 1999; *speciesLink network*, 2021). Similarly, aeginetin has only been reported in plants of the Buchnerae clade, including *Aeginetia* (subclade H, highly related to subclade I), *Centranthera* (Eschenmoser *et al*., 1982; Zhang *et al*., 2019; Nickrent, 2020), and now *Escobedia*. It is possible, therefore, that the production of these apocarotenoids, and particularly the accumulation of azafrin, might be play a biological function related to root parasitism in this specific group of plants.

### Analysis of the Escobedia root transcriptome allows the identification of candidate genes for carotenoid biosynthesis

To identify tentative genes involved in the synthesis of azafrin in *Escobedia*, we extracted RNA from a pool of orange roots collected from different plants in a natural population and used it for RNA-seq analysis. Following adapter removal, trimming, and quality filtering with *fastp*, 76 million 2×150 bp paired-end reads (~ 10.8 Gb of sequence data) were used for *de novo* transcriptome assembly using *Trinity*. Summary statistics of the final assembly include, among others, a total of 115,882 transcripts (65,701 ‘genes’) ranging between 200 and 12,858 bp in length, a mean and N50 sequence length of 1,111 and 1,798 bp, a GC-content of 0.43, and a 25.5% of total transcripts deemed as lowly or not expressed (i.e., FPKM < 0.5) (Supporting Information Fig. S1a). Additionally, 61,325 and 48,576 transcripts showed evidence-supported coding sequences predicted using *TransDecoder* and *EvidentialGene* pipelines, respectively. Transcripts containing predicted CDS by *TransDecoder* revealed that many were complete (i.e., full-length and containing both start and stop codons) and predominantly being >1kb in length (Supporting Information Fig. S1b). BUSCO assessments with the embryophyta lineage database also indicated very high predicted completeness of the full transcriptome assembly and reduced CDS-only set (i.e., 93.7% - 94.2% of 1,375 BUSCOs evaluated) (Supporting Information Fig. S1c). These BUSCO scores rival those observed in many sequenced plant genomes with high-quality gene models (Veeckman *et al*., 2016).

Functional annotation according to MapMan BIN v4 categories, matching Pfam domain of predicted peptides, or shared homology with plant Uniprot database sequences, revealed 51,302, 51,578, and 66,335 transcripts (44 – 57% of total transcripts), respectively (Supporting Information Table S2). Notably, we observed a greater representation of transcripts associated to MapMan BIN15_RNA biosynthesis, BIN18_Protein modification, BIN19_Protein homeostasis, BIN24_Solute transport, and BIN50_Enzyme classification, among others (Supporting Information Fig. S1d). More relevant for azafrin biosynthesis, BIN categories related to biosynthetic pathways of carotenoids (BIN9.1.6.1) and apocarotenoids (BIN9.1.6.3), among others, were also successfully assigned to transcripts (Supporting Information Fig. S1d). Additionally, 45,339 and 68,727 transcripts present in *Escobedia* roots shared orthology with *Arabidopsis* and *Centranthera* genes, respectively (Supporting Information Table S2). Together, the assembly statistics, gene completeness scores, and number of functional categories and domain assignments to transcripts suggest a reasonably high assembly quality of the pooled root transcriptomes.

According to our annotation pipelines, we identified thirty-four genes (30 predicted complete and 4 partial sequences) potentially involved in carotenoid and apocarotenoid pathways that were expressed in *Escobedia* roots (Fig. 2; Supporting Information Table S3). The first committed step of the carotenoid pathway is the production of phytoene from GGPP catalysed by PSY (Fig. 2a). PSY, the main flux-controlling enzyme of the plant carotenoid pathway (Fraser *et al*., 2002; Rodriguez-Concepcion *et al*., 2018), is usually encoded by small gene families encoding distinct isoforms associated with organ- or tissue-specific production of carotenoids. For example, tomato PSY1 is essential for fruit carotenoid production during ripening, while PSY2 is preferentially found in photosynthetic tissues, and PSY3 functions in the root. The root-associated PSY3 isoforms from dicots form a widespread phylogenetic clade found to participate in arbuscular mycorrhiza (AM) interactions, whereas those from monocots form a different clade and are involved in ABA formation (Stauder *et al*., 2018). Two PSY isoforms were found to be expressed in *Escobedia* roots, namely PSYa and PSYb (Fig. 2a). The phylogenetic comparison of the *Escobedia* PSYa and PSYb isoforms with PSY sequences from *Centranthera* and two other root hemiparasitic species (*Phtheirospermum japonicum* and *Striga asiatica*,) together with well-characterized PSY sequences from monocots (maize, rice) and dicots (tomato, carrot, alfalfa, *Arabidopsis*) led to their classification in a sub-clade of only root hemiparasitic PSY sequences with a 100% of posterior probability (Fig. 3). This subclade, however, was separated from the clades harbouring root-associated PSY3 sequences from dicots or monocots (Fig. 3). Similar to *Escobedia*, two genes encoding PSY and belonging to the same subclade were found in *Centranthera* (Fig. 3). It is interesting to note that the *Centranthera* gene encoding PSYa was more actively expressed in leaves and stems than in roots whereas higher levels of transcripts encoding PSYb were found in azafrin-producing roots compared to leaves and stems (Fig. 2a) (Zhang *et al*., 2019). These results suggest that some non-PSY3 isoforms of PSY might have a prominent role in the roots of at least some hemiparasitic plants. In the case of *Escobedia*, genes encoding both PSYa and PSYb isoforms are expressed at similarly high levels in roots, suggesting that azafrin production may require an active metabolic flux into the carotenoid pathway.

**Fig. 2.**
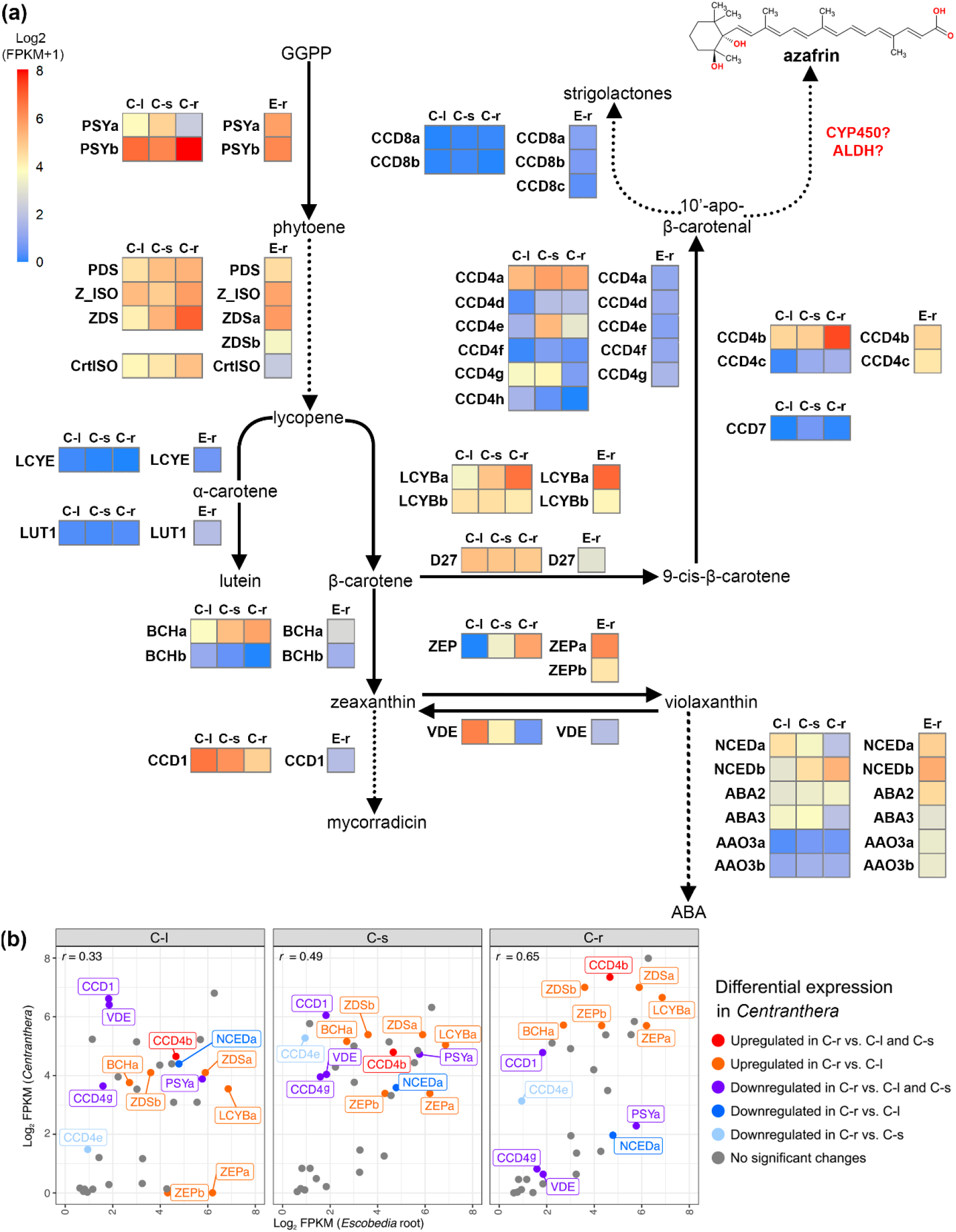
Candidate genes of the azafrin biosynthetic pathway in *Escobedia* and *Centranthera*. **(a)** Proposed pathway and abundance of enzyme-encoding transcripts in *Escobedia* roots (E-r) and *Centranthera* leaf (C-l), stem (C-s) and root (C-r) tissues. Dotted lines represent multiple steps. Colours represent transcript abundance based on RNA-seq analyses (acronyms and FPKM values are listed in Supporting Information Table S3). Transcripts for the enzymes indicated in red were not identified in the root transcriptome. **(b)** Spearman correlation of gene expression values between *Escobedia* roots and the indicated *Centranthera* tissues. Scale corresponds to log2 transformed values (FPKM+1). Data for *Centranthera* were retrieved from Zhang et al. (2019) and reanalysed.

**Fig. 3.**
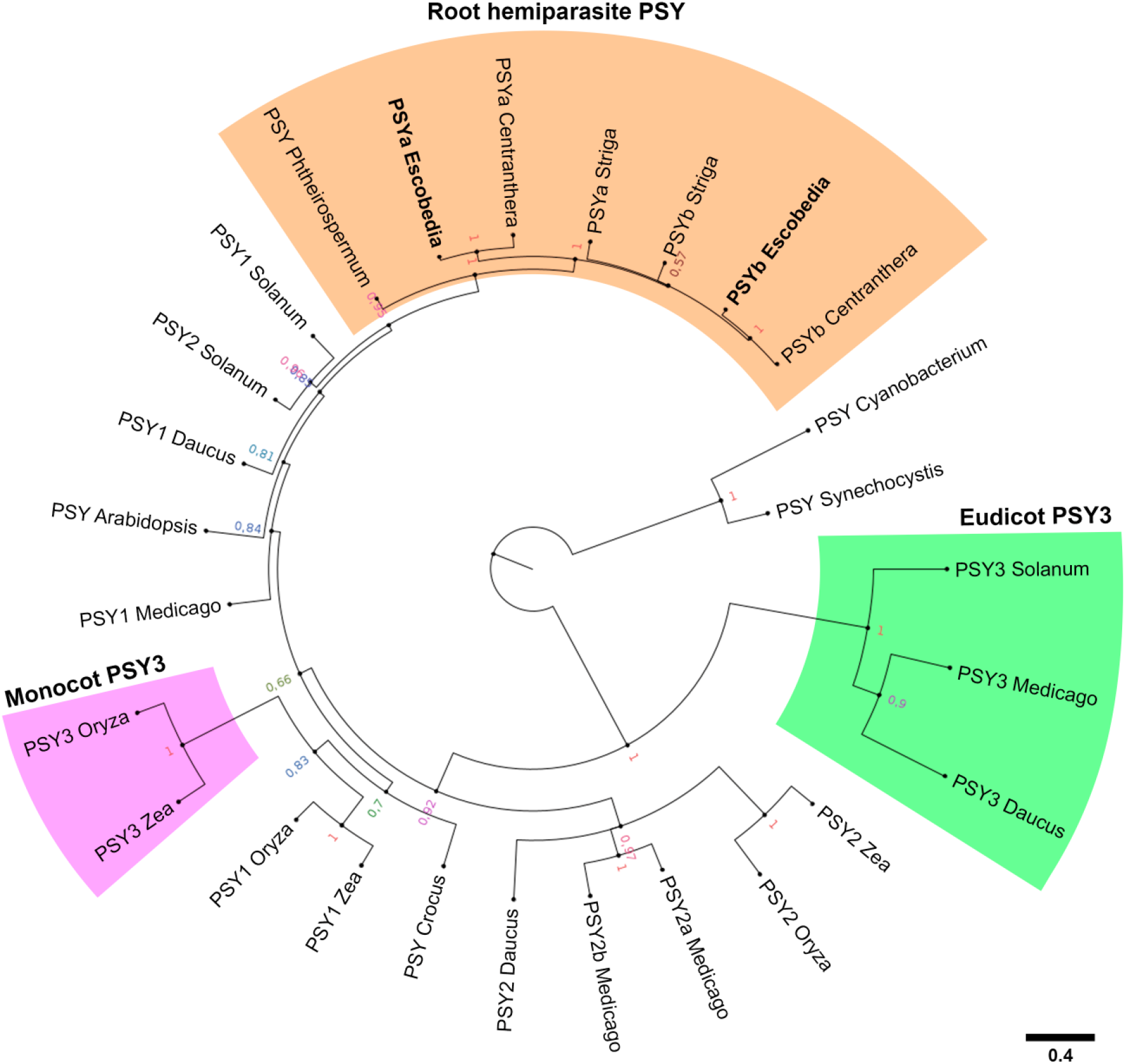
Phylogenetic analysis of the phytoene synthase (PSY) family in several plants. The computed Bayesian tree includes twenty-five angiosperm species and two bacteria as outgroups. The total number of generations is 200,000. The average standard deviation of split sequences is 0.019. Distinctively separate clades are indicated with colours. *Escobedia* sequences are in bold. Gene accessions are listed in Supporting Information Table S4.

In the next section of the carotenoid pathway (i.e., from phytoene to lycopene), transcripts corresponding to single genes were found for phytoene desaturase (PDS), ζ-carotene isomerase (Z-ISO), and carotenoid isomerase (CRTISO), whereas two *Escobedia* genes were expressed encoding ζ-carotene desaturase (ZDS) (Fig. 2a). Interestingly, *Arabidopsis* mutants defective in ZDS have been reported to produce an apocarotenoid signal that negatively impacts plastid and leaf development (Avendaño-Vázquez *et al*., 2014; Escobar-Tovar *et al*., 2021). Thus, the relatively high expression levels of the two ZDS-encoding genes detected in *Escobedia* roots (Fig. 2a) might result in high ZDS activity and hence prevent the formation of the apocarotenoid signal generated in ZDS-defective mutants. After lycopene, the carotenoid pathway branches out (Fig. 2a). Cyclization of the two ends of the linear lycopene molecule to produce one β and one ε ring generates α-carotene. These reactions are catalysed by lycopene β and ε cyclases (LCYB and LCYE, respectively). By contrast, β-carotene is synthesized from lycopene when only β rings are formed by LCYB enzymes. Transcripts encoding LCYE were found at much lower levels than those encoding the two LCYB isoforms expressed in *Escobedia* and *Centranthera* roots, LCYBa and LCYBb (Fig. 2a), suggesting a higher flux through the β, β branch compared to the β, ε branch. In agreement with this conclusion, transcripts for the LUT1 enzyme, which catalyses the hydroxylation of ε rings to produce lutein (the most abundant carotenoid in green photosynthetic tissues), are much less abundant than those for β-ring hydroxylases (BCHa and BCHb). Based on the higher abundance of transcripts for BCHa in both *Escobedia* and *Centranthera*, this might be the main BCH isoform transforming β-carotene into zeaxanthin (Fig. 2a). Then, two highly expressed zeaxanthin epoxidase isoforms (ZEPa and ZEPb) in *Escobedia* and a single ZEP-encoding gene in *Centranthera* produce violaxanthin in roots. The very low levels of transcripts found for violaxanthin epoxidase (VDE) suggest that the main metabolic flux is from zeaxanthin to violaxanthin in *Escobedia* and *Centranthera* roots.

### Azafrin production in Escobedia roots might rely on CCD4 rather than CCD7 enzymes

Presence of transcripts encoding NCED and other ABA biosynthetic enzymes such as ABA2, ABA3, AAO3 (Fig. 2a), suggest that violaxanthin can be converted into ABA in azafrin-producing *Escobedia* and *Centranthera* roots. Besides ABA, other apocarotenoids are formed by the activity of CCD enzymes. Among them, SL and azafrin production are proposed to share the first isomerization step from β-carotene (Fig. 2a). Transcripts encoding D27 were found in azafrin-accumulating roots, supporting the conclusion that β-carotene can be isomerized to 9-*cis*-β-carotene in this tissue to allow the production of SL or/and azafrin (Fig. 2a). Both apocarotenoids were also proposed to share the next step of the pathway, i.e. the cleavage of C_40_ 9-*cis*-β-carotene by CCD7 to produce C_27_ 10’-apo-β-carotenal (Zhang *et al*., 2019). In-depth phylogenetic analysis of the CCD family from *Escobedia, Centranthera* and several other plants, including hemiparasitic species, showed four clades designated as CCD1 (paraphyletic), CCD4, CCD7, and CCD8 (Fig. 4). The only gene encoding CCD1 was found to be poorly expressed in *Escobedia* roots, whereas in *Centranthera* it was less expressed in roots than in other tissues such as leaves and stems (Fig. 2a). The CCD4 clade contained seven CCD4 sequences from *Escobedia* (CCD4a to g), of which CCD4b and CCD4c were most highly expressed in *Escobedia* roots (Fig. 2a). Of the eight CCD4 sequences found in *Centranthera* (CCD4a to h), only CCD4d was predominantly expressed in the azafrin-producing roots (Fig. 2).

**Fig. 4.**
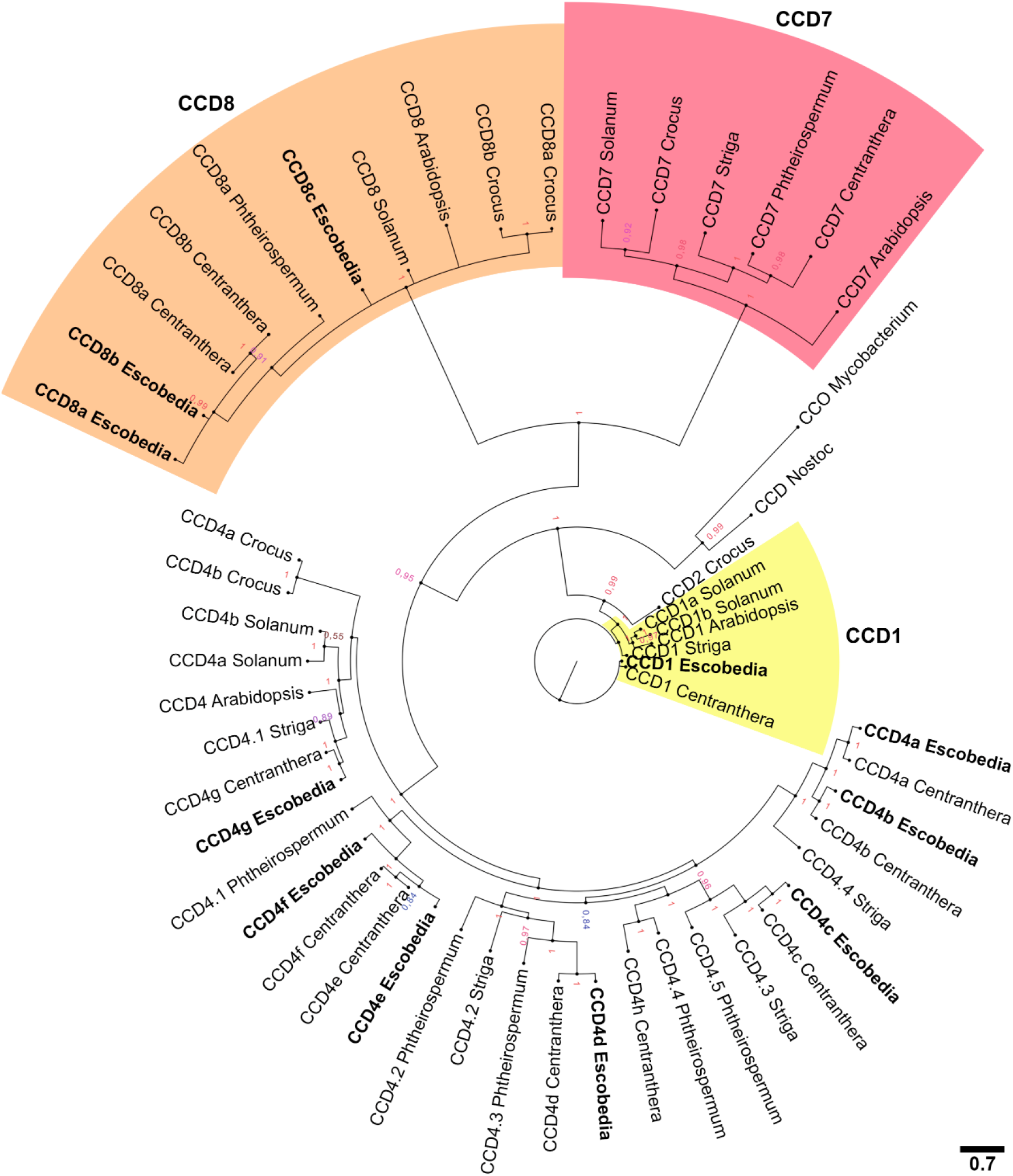
Phylogenetic analysis of the carotenoid cleavage dioxygenase (CCD) family in several plants. The computed Bayesian tree includes fifty-one sequences from seven angiosperms and two bacteria as outgroups. The total number of generations is 185,000. The average standard deviation of split sequences is 0.02. Distinctively separate clades are indicated with colours. *Escobedia* are in bold. *Escobedia grandiflora* sequences are in bold text. Asterisks indicate incomplete amino acid sequences. Gene accessions are listed in Supporting Information Table S4.

A striking conclusion of our phylogenetic analysis is that no sequences of *Escobedia* were present in the CCD7 clade, which included sequences of *Centranthera* and other hemiparasitic species as well as typical CCD7 enzymes from *Arabidopsis* and tomato. We hence speculated that *Escobedia* roots CCD4b or/and CCD4c might catalyse the same C9-C10 cleavage reaction that CCD7 enzymes perform in other plants or tissues. The CCD4 subfamily is probably the most variable group of CCD enzymes. They are encoded by several genes in many species, resulting in isoforms that often differ in their expression profile and substrate selectivity (Hou et al., 2016). In general, CCD4 enzymes have broad substrate specificity, and many of them appear to have a role in carotenoid catabolism, particularly in carotenoid-sink tissues such as flowers, fruits, seeds, and roots (Walter *et al*., 2010; Rubio-Moraga *et al*., 2014). They usually cleave carotenoids (notably β-carotene) at the C9–C10/C9’-C-10’ double bond, but the *Arabidopsis* CCD4 enzyme also catalyses the C9–C10 cleavage of β, β xanthophylls such as zeaxanthin while other CCD4 enzymes cleave asymmetrically at the C7– C8/C7’-C-8’ double bond (Rubio *et al*., 2008; Huang *et al*., 2009; Ma *et al*., 2013; Lätari *et al*., 2015; Bruno *et al*., 2016). Interestingly, *Arabidopsis* CCD4 has been shown to catalyse the cleavage of β-carotene to all-*trans*-10’-apo-β-carotenal and, at a much lower efficiency, of 9-*cis*-β-carotene to the SL precursor 9-*cis*-10’-apo-β-carotenal (Bruno *et al*., 2016). It can thus be suggested that *Escobedia* CCD4b or/and CCD4c isoforms might also catalyse these reactions to deliver precursors for SL and azafrin biosynthesis, hence making the participation of a CCD7 enzyme unnecessary. Interestingly, in the roots of *Centranthera* transcripts encoding CCD7 were expressed at much lower levels than those encoding D27 (Fig. 2a), whereas CCD4b was much more highly expressed than CCD7 (Fig. 2a) (Zhang *et al*., 2019). It is therefore possible that CCD4 isoforms might produce the precursors for SL and azafrin in both *Escobedia* and *Centranthera* roots. Further supporting this conclusion, our analysis indicated that *Centranthera CCD4b* is the only gene that shows root-specific differential upregulation compared to stem and leaf tissues among all other azafrin-related *Escobedia* orthologs (Fig. 2b; Supporting Information Table S3).

To complement the RNA-seq analysis, transcript levels encoding CCD4b and CCD4c were measured in roots of *Escobedia* plants grown either with or without hosts (Fig. 5). Previous studies observed that the roots of *Escobedia* were colourless in the initial developmental stages (i.e., before parasitizing a host), while the orange pigment became most visible after haustoria penetration in the host root (Cardona-Medina *et al*., 2019). HPLC analyses confirmed that azafrin levels were substantially increased in *Escobedia* roots when growing in the presence of host plants compared to those grown in their absence (Fig. 5). The level of transcripts encoding CCD4c were similar regardless of the presence of a host, whereas those encoding CCD4b were present at much higher levels in azafrin-producing roots (Fig. 5). Together, these results suggest that CCD4b might be the main enzyme participating in the production of azafrin.

**Fig. 5.**
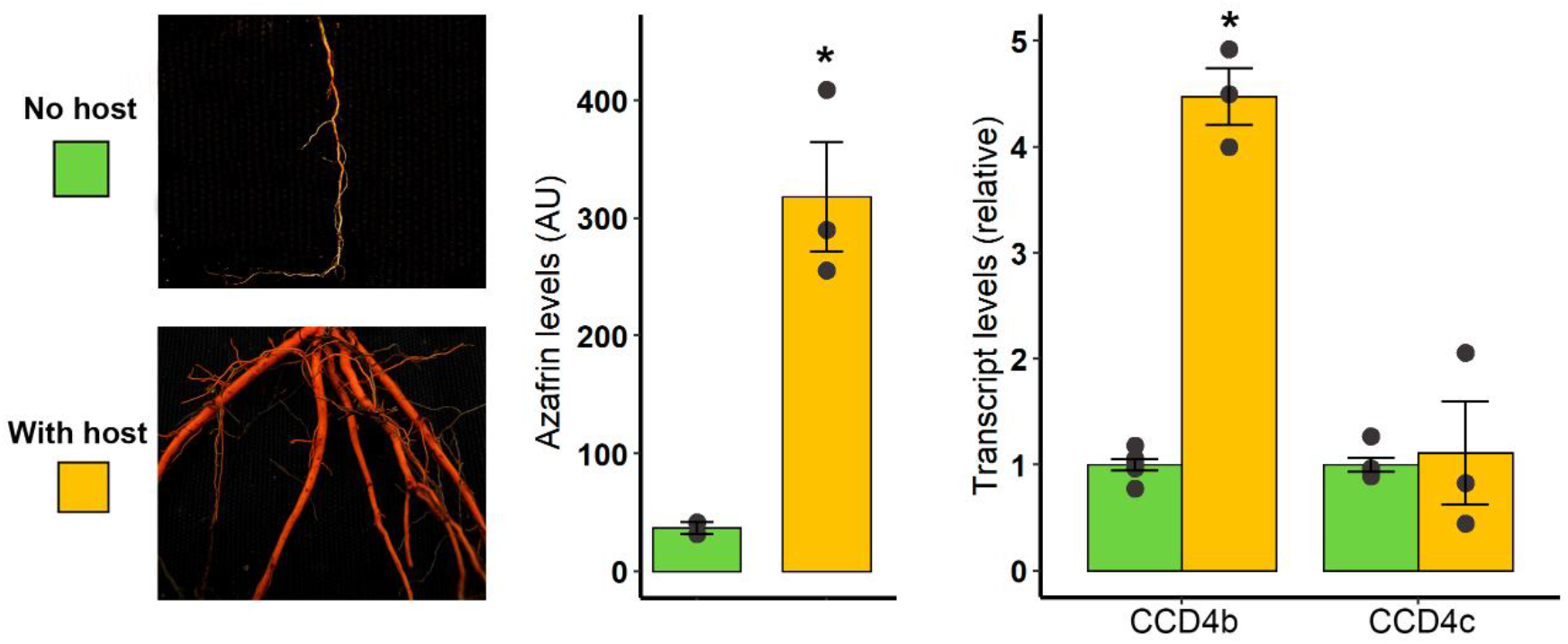
Levels of azafrin and transcripts encoding CCD4 homologs in roots of *Escobedia* plants grown with (orange) and without a host (green). Asterisks represent significant differences by Tukey test (*P*<0.05). Error bars represent standard error of the means (n=3).

C_27_ 10’-apo-β-carotenal is the substrate of CCD8 enzymes in the next step of the SL biosynthesis pathway (Fig. 2a). *Escobedia* and *Centranthera* genes encoding homologs for CCD8 were found to be expressed at much lower levels than the one for D27 in roots (Fig. 2a). Based on these data, we hypothesise that the pathway for producing strigolactones (via CCD8) is not very active in azafrin-producing *Escobedia* and *Centranthera* roots. This might not be a general trend in parasitic plants. For example, the tubercle of the holoparasitic plant *Phelipanche aegyptiaca* showed a high expression of D27, CCD7, and CCD8 when parasitizing the host roots (Emran *et al*., 2020). However, mycorrhizal plants that produce high levels of apocarotenoids reduce their production and secretion of SL, likely because abundant apocarotenoid production in AM-colonized roots might generate a metabolic sink and successfully compete for SL precursors (Walter *et al*., 2010). Similarly, the presumably low flux towards SL in azafrin-producing *Escobedia* and *Centranthera* roots might be related to the requirement of very high levels of common precursors to support the massive production of azafrin in the roots of these hemiparasitic plants. Strikingly, we were unable to detect any carotenoid species in *Escobedia* roots, suggesting that the carotenoid pathway in this organ is mainly directed to provide substrates for apocarotenoid production, and no intermediates are accumulated. Similarly, no detectable amounts of carotenoids were found in AM roots of all plants investigated despite the required high flux through the carotenoid pathway (Fester *et al*., 2002).

### Azafrin accumulates in the apoplast of the Escobedia root cortex

An anatomical analysis was next performed to investigate where azafrin accumulated in the roots of *Escobedia* plants collected in the wild. The light microscopy visualization of the internal structure of the root revealed that the orange pigment corresponding to azafrin was not located in plastids (the site where all plant carotenoids are made) or in the cell cytosol (where the synthesis of many apocarotenoids is completed) but accumulated in the intercellular spaces (i.e., apoplast) of the root cortex (Fig. 6a). Confocal laser scanning microscopy analyses based on the autofluorescence produced by the c.b.d. system present in the polyene chain of carotenoids and apocarotenoids of sufficient length such as C_27_ azafrin (D’Andrea *et al*., 2014) confirmed that the orange pigmentation detected by light microscopy was due to the presence of azafrin as both autofluorescence and color signals overlapped (Fig. 6b; Supporting Information Fig. S2). Light and confocal microscopy of carrot (*Daucus carota*) roots also showed an overlap of color and autofluorescence signals, but in this case they were both detected inside plastids (i.e., chromoplasts) as they correspond to carotenes (β-carotene and, to a lower extent, α-carotene) instead of their cleavage products (Supporting Information Fig. S2). By contrast, azafrin was virtually absent from the large starch-filled plastids (i.e., amyloplasts) present in *Escobedia* roots (Fig. 6; Supporting Information Fig. S2).

**Fig. 6.**
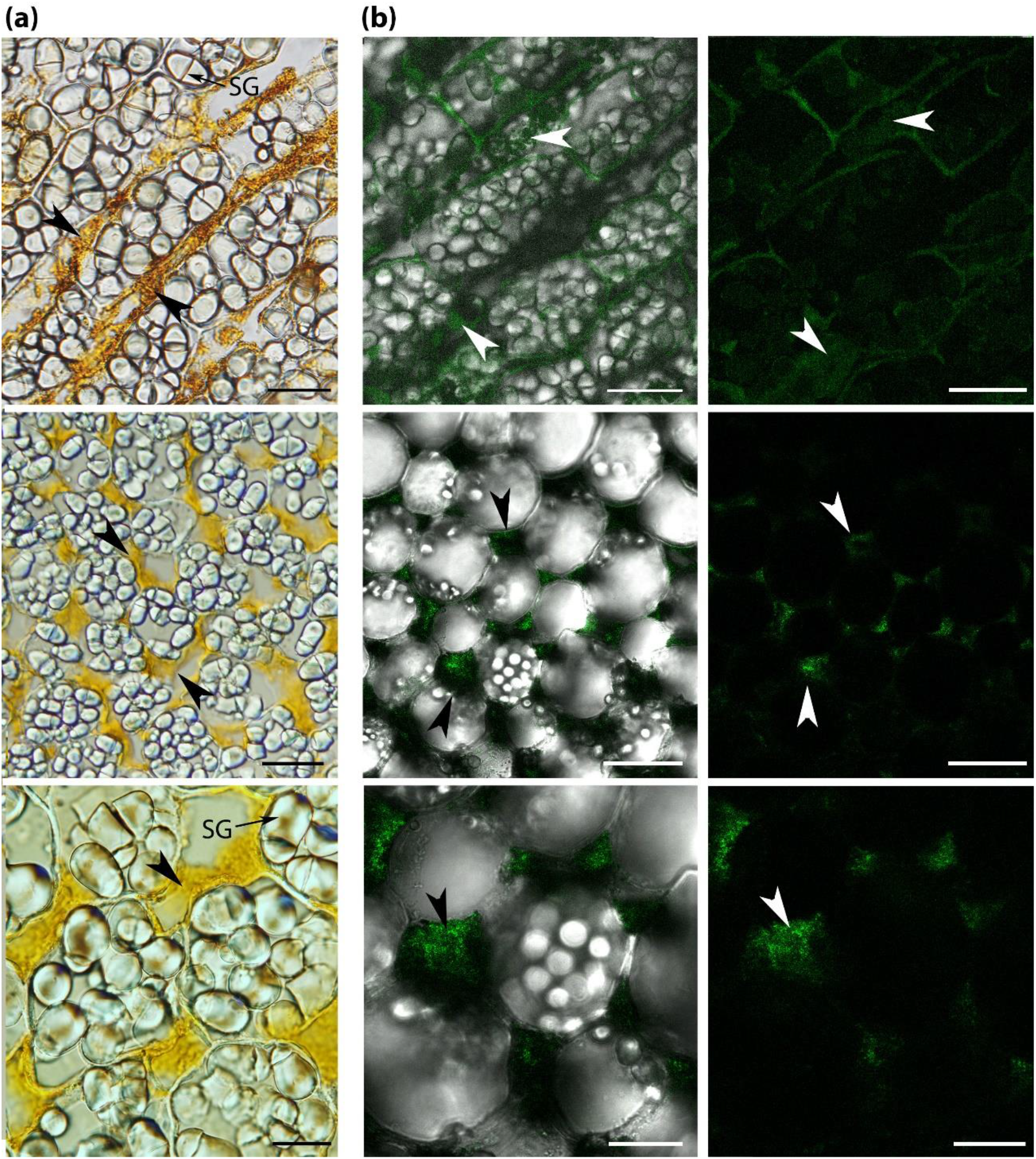
Azafrin accumulates in the apoplastic space of the *Escobedia* root cortex. **(a)** Micrographs under light microscopy show azafrin as an orange pigment in the intercellular spaces (apoplast) of the cortex (arrowheads). **(b)** Micrographs under confocal microscopy show azafrin as green autofluorescence (arrowheads). Fluorescence images (right panels) are shown next to the corresponding merged micrographs of fluorescence and bright field images of the same field. Images show representative images of root longitudinal sections (upper panels) and cross-sections (central and lower panels). SG, starch grains. Scale bars: 50 μm (upper and central panels), 20 μm (lower panels).

Carotenoids are synthesized and accumulated in different plastid types, including amyloplasts (Horner *et al*., 2007; Sun *et al*., 2018; Rodriguez-Concepcion *et al*., 2018). Studies of apocarotenoid formation in mycorrhizal roots have led to conclude that the C_27_ products of CCD4 or/and CCD7 activities are exported from the plastid and used in the cytosol as substrates of CCD1 enzymes that convert them into downstream products such as C_13_ (colorless) blumenols and C_14_ (yellow) mycorrhadicins (Walter *et al*., 2010; Fiorilli *et al*., 2019; Moreno *et al*., 2021). Similarly, the potential C_27_ product of CCD4b activity in the amyloplasts of *Escobedia* roots might be exported from the plastids to the cytosol. Accumulation of C_27_ apocarotenoids is uncommon in nature, probably because CCD1 enzymes normally degrade them (Floss *et al*., 2008; Walter *et al*., 2010). The low expression level of the only gene encoding CCD1 in *Escobedia* roots (Fig. 2a) suggests that most of this C_27_ intermediate might remain available for other cytosolic enzymes to transform it into downstream products. The differences between azafrin and 10’-apo-β-carotenal are one terminal carboxyl group and two hydroxyl groups in the cyclohexane skeleton (Zhang *et al*., 2019). The activity of cytosolic aldehyde dehydrogenase and cytochrome P450 monooxygenase enzymes could transform the aldehyde group of 10’-apo-β-carotenal into carboxylic acid and insert oxygen atoms, respectively, eventually improving hydrophilicity (Fig. 2a). Water-soluble azafrin might then be released from the root cells and accumulate in the apoplast. Based on the putative role reported for the accumulation of colored apocarotenoids in AM-inoculated roots, it is possible that azafrin might participate in the interaction with the rhizosphere, e.g. by providing protection from oxidative damage caused by biotic or abiotic stresses (Strack & Fester, 2006).

### Evidence for a possible role for azafrin in the parasitization process

Following germination, the root of *Escobedia* is colourless and seedlings grow very slowly until a host is parasitized, which involves both the formation of specialized organs (haustoria) to penetrate the host root and the pigmentation of the root, i.e. the accumulation of azafrin (Cardona & Muriel, 2015; Cardona-Medina *et al*., 2019). The haustorium is an organ characteristic of parasitic plants that have evolved in multiple independent angiosperms. It contains structures for mechanical attachment to the host root and vascular connections that involve the differentiation of various specialized cell types (Teixeira-Costa, 2021). A detailed exploration of this structure in *Escobedia* showed numerous tubular haustorial hairs associated with the orange swelling periphery (Supporting Information Fig. S3). These hairs participate in securing the haustorium to the epidermis of the host root and facilitate the contact and penetration into host tissues (Heide-Jorgensen & Kuijt, 1995; Cui *et al*., 2016). The haustorial opening, i.e., the area that makes contact with the host root, was observed in the haustorium apex (Supporting Information Fig. S3a). Longitudinal sections of the *Escobedia* haustorium attached to host roots revealed a complex internal structure, presenting four recognizable regions (Supporting Information Fig. S3b): haustorial base, vascular tissue, hyaline body, and intrusive cells (endophyte). The haustorial base connects the parasitic root with the haustorium, morphologically similar to root tissue. The vascular tissue comprised of provascular cells and tracheary elements (haustorium xylem) is arranged perpendicularly to the haustorial base and towards the host root xylem. Intrusive cells of the haustorium penetrate and advance inside the host root, constituting the endophyte. Tracheary elements of the haustorium were observed inside the host’s metaxylem, indicating parasitism success in the host root (Supporting Information Fig. S3b). Interestingly, we noticed that every haustorium attached to host roots contained orange pigment depositions corresponding to azafrin in the region directly contacting the host root interface (Fig. 7; Supporting Information Fig. S3b).

**Fig. 7.**
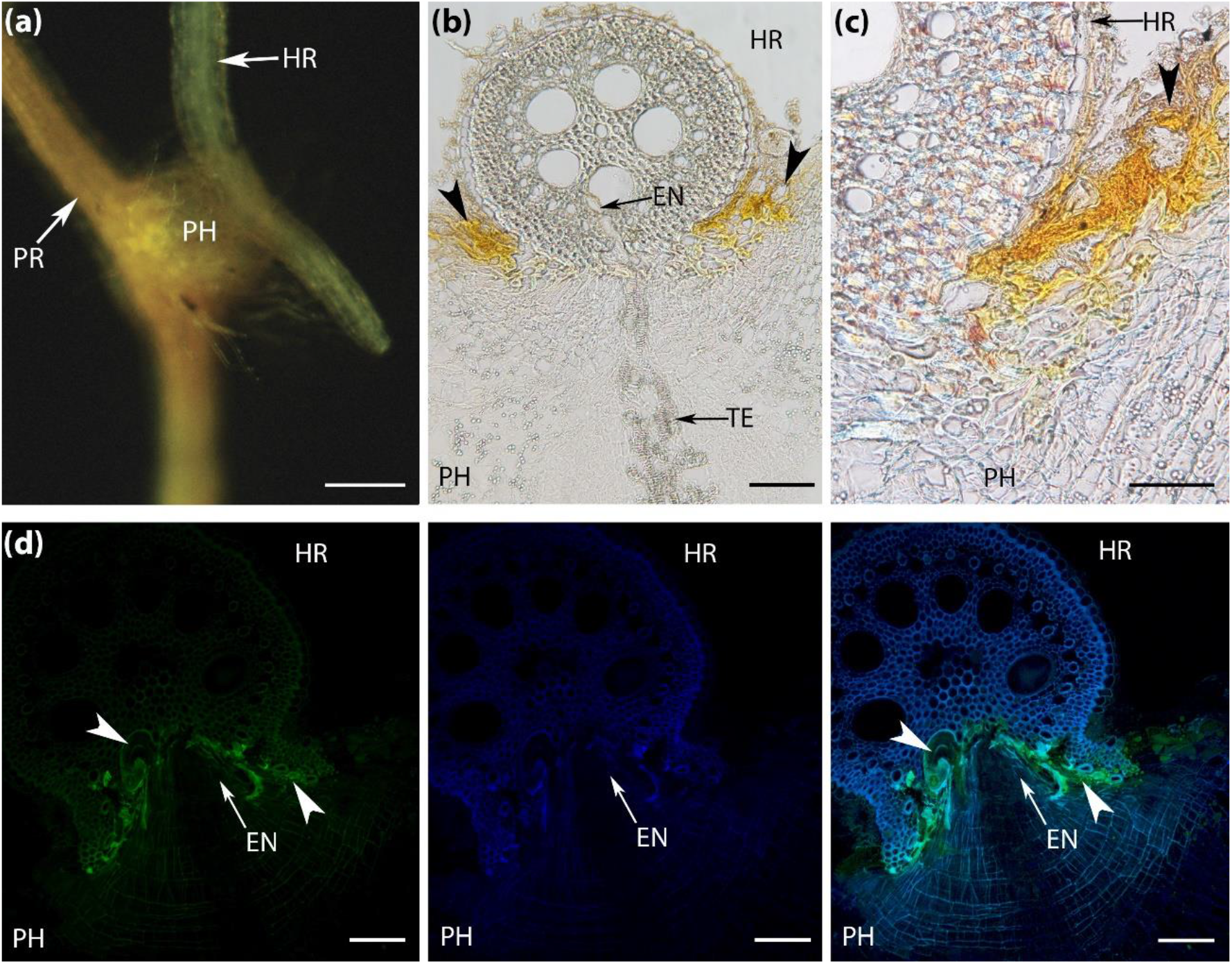
Azafrin accumulates in the haustorium-host root interphase. **(a)** External view of *Escobedia* haustorium attached to a host (*Pennisetum purpureum*) root. **(b)** Representative light microscopy picture of the haustorium-host root interface showing an accumulation of the orange pigment azafrin (arrowhead). **(c)** Detail of azafrin pigment accumulation in the haustorium-host root interface (arrowhead). **(d)** Representative confocal microscopy images showing the localization of azafrin-associated fluorescence in the haustorium-host root interface. From left to right: green autofluorescence corresponding to azafrin (arrowhead); blue autofluorescence corresponding to cell walls; and merged green and blue autofluorescence. HR, host root; EN, endophyte; PH, parasite haustorium, PR, parasite root; TE, tracheary elements. Scale bars: 200 μm (a), 100 μm (b, d), 50 μm (c).

The reason why azafrin accumulates in the haustorium-host root interface is still unknown. We propose that azafrin might inhibit the host defence responses during the penetration of the haustorium inside the roots. Haustorium penetration in host roots involves enzymatic secretion and mechanical pressure (by haustorial hairs and cellular division) that degrade and disrupt the host cells walls (Heide-Jorgensen & Kuijt, 1995; Hood *et al*., 1998; Losner-Goshen *et al*., 1998). This process causes a wound in the host root that allows the entry of haustorium intrusive cells into the host vascular system. Host roots quickly respond to the wound by activating the production of reactive oxygen species (ROS), which can activate programmed cell death (PCD) and trigger plant defence responses (Minibayeva *et al*., 2009; Tripathy & Oelmüller, 2012), including the generation of phenolic compounds and callose deposition, induction of immunity-related genes, and deposition of lignin and suberin to avoid the advance of parasitization (Hiraga *et al*., 2001; Minibayeva *et al*., 2009; Saucet & Shirasu, 2016). Evidence of necrosis involving ROS was found in resistant hosts during unsuccessful penetration by the haustorium of *Orobanche cumana* (Letousey *et al*., 2007). Thus, the presence of azafrin in the haustorium might inhibit host PCD and defense responses by eliminating extracellular ROS as a strategy to facilitate parasitization (Mor *et al*., 2008). Inactivation of defence responses has also been observed in biotrophic pathogens during the parasitism of host plants, due to its need to proliferate in living host cells (Siddique *et al*., 2014). Likewise, endophytic fungi can produce antioxidants to circumvent damage by ROS during beneficial interaction with the host plant (Hamilton *et al*., 2012). Together, we speculate that azafrin accumulation in the haustorium-host root interface might also play an antioxidant role to counteract the ROS-related host defence responses, hence allowing the parasitism to succeed. Further work should experimentally address this and other unanswered questions about the role of azafrin and other apocarotenoids in the parasitization strategy of *Escobedia*. The information would be vital to eventually achieve the domestication and cultivation and hence exploitation of this important medicinal plant.

## Acknowledgements

We are grateful to Xiaodong Zhang and Wensheng Qin for providing *Centranthera* sequences. We thank María Rosa Rodríguez-Goberna (CRAG), Montserrat Amenós (CRAG) and Eliana Medeiros (LCME-UFSC) for technical support. We thank Paula Astolfi for the initial review of this manuscript. We thank to Instituto Chico Mendes de Conservação da Biodiversidade (ICMBio), the national major conservation agency, and Fundação Municipal do Meio Ambiente (FLORAM), the Florianópolis environmental agency, who issued the authorizations n°63251-8 and n°020/18-DEPUC for the fieldwork.

ECM was supported by CAPES, The Brazilian Agency for Higher Education, and the project PrInt CAPES-UFSC. RON is supported by National Council for Scientific and Technological Development, Brazil (CNPq) (Proj. 303902/2017-5). JTM is supported by Grants PGC2018-099449-A-I00 and RYC-2017-23645 from MCIN/AEI/10.13039/501100011033 and “ERDF A way of making Europe”. MRC lab is funded by grants PID2020-115810GB-I00 and PCI2021-121941 (funded by MCIN/AEI/10.13039/501100011033 and the European Union NextGeneration EU/PRTR and PRIMA-UToPIQ), 202040E299 (funded by Consejo Superior de Investigaciones Científicas, CSIC), and PROMETEU/2021/056 (funded by Generalitat Valenciana).

## Author Contributions

ECM, RON and MRC designed the research; ECM collected plant material and performed the experiments; ECM and DHM carried out azafrin analysis; AP, JTM and DW conducted all bioinformatic analyses; ECM, MS, RON, MRC performed microscopical analyses; ECM and MRC wrote the paper. All authors contributed to interpretations and revisions of the manuscripts.

## Data availability

RNA-Seq original sequence data can be found in the Sequence Read Archive (SRA) database of the NCBI under the Bioproject accession PRJNA798758.

## Supporting Information

**Methods S1.** Azafrin detection and quantification.

**Methods S2.** Transcriptome analysis and functional annotation

**Methods S3.** RT-qPCR.

**Methods S4.** Structural and anatomical analyses.

**Fig. S1** *De novo* transcriptome assembly from *Escobedia* roots.

**Fig. S2.** Carotenoid and azafrin distribution in carrot and *Escobedia* roots.

**Fig S3**. Overview of the *Escobedia* haustorium.

**Data S1.** Nucleotide sequences of *Escobedia* PSY and CCD homologs.

**Table S1.** Primers used for RT-qPCR.

**Table S2.** *De novo* transcriptome assembly using Trinity.

**Table S3.** Accessions and FPKM values of *Escobedia* and *Centranthera* transcripts involved in azafrin biosynthesis.

**Table S4.** PSY and CCD accessions used in phylogenetic analyses.

