## Supplemental methods and figures for "Biosynthesis and apoplast accumulation of the apocarotenoid pigment azafrin in parasitizing roots of *Escobedia grandiflora*"

The following Supporting Information is available for this article:

**Methods S1.** Azafrin detection and quantification.

**Methods S2.** Transcriptome analysis and functional annotation

**Methods S3.** RT-qPCR.

**Methods S4.** Structural and anatomical analyses.

**Fig. S1** *De novo* transcriptome assembly from *Escobedia* roots.

**Fig. S2** Carotenoid and azafrin distribution in carrot and *Escobedia* roots

**Fig. S3** Overview of the *Escobedia* haustorium.

**Data S1** Nucleotide sequences of *Escobedia* PSY and CCD homologs. (see Fasta file)

**Table S1.** Primers used for RT-qPCR. (see Excel file)

**Table S2.** *de novo* transcriptome assembly of *Escobedia grandiflora* roots using Trinity. (see Excel file)

**Table S3.** Code, identification and FPKM values used in the proposed pathway for azafrin biosynthesis in *Escobedia grandiflora* and the orthologs of *Centranthera grandiflora* leaf, stem and roots. (see Excel file)

**Table S4.** Gene code and IDs of amino acid sequences of PSY and CCDs used in phylogenetical analysis. (see Excel file)

#### Supporting Information - Methods

**Methods S1. Azafrin identification and quantification.**

For pigment extraction, 10 mg of freeze-dried root powder was mixed with 10 ml of methanol and mixed. Then, methanol was evaporated in a rotavapor (at less than 30ºC) and the remaining extract was resuspended in 0.5 ml of methanol for HPLC-MS(APCI) analysis using a Dionex Ultimate 3000RS U-HPLC (Thermo Fisher Scientific) coupled in series with a diode array detector (DAD) and a micrOTOF-QII high resolution time-of-flight mass spectrometer (UHR-TOF) with qQ-TOF geometry (Bruker Daltonics) and fitted with an APCI source. A flow-split of the eluent from the DAD detector was set up in order to allow a 0.4 mL/min flow rate directly into the mass spectrometer (connected in series after the DAD detector). The instrument control was performed using Bruker Daltonics Hystar 3.2 software and data evaluation was performed with the Bruker Daltonics DataAnalysis 4.0 software. The chromatographic conditions were identical to those for HPLC-DAD analysis (see below). Briefly, separation was carried out by a binary-gradient elution using an initial composition of 75 % acetone and 25 % deionized water (both containing 0.1 % formic acid), which was increased linearly to 95 % acetone in 10 min, then hold for 7 min and raised to 100 % in 3 min, and maintained constant for 10 min. Initial conditions were reached in 5 min. The separation was performed in a reversed-phase C18 (20 mm x 4.6 mm i.d., 3 µm, Mediterranea SEA18; Teknokroma) fitted with a guard column of the same material. The temperature of column was kept at 25 ºC. An injection volume of 20 µL and a flow rate of 1 mL/min were used. UV-visible detection was performed at 450 nm and the online spectra were acquired in the 350–700 nm wavelength range. The MS parameters were set as follows: positive mode; current corona, 4000 nA; source (vaporizer) temperature, 350 °C; drying gas, N2; gas temperature, 250 °C; gas flow, 4 L/min; nebulizer pressure, 60 psi; scan range of m/z 50–1000. For azafrin quantification, 1 mg of lyophilized root powder was mixed with 1 ml of methanol. After mixing by agitation for 1 min and centrifugation at 1300 rpm for 5 min at 4°C in a microfuge, the upper phase was collected and filtered through a 0.2 µm microfilter. For HPLC-DAD, 10 µL of each sample were injected onto an Agilent Technologies 1290 Infinity system and azafrin was monitored at 450 nm using the Agilent ChemStation software.

**Methods S2. Transcriptome analysis and functional annotation**

Surviving paired-end reads were aligned against the reference assembly using *bowtie2* (Langmead & Salzberg, 2012) using the local read alignment mode (--*local*). Read summarization was performed using *FeatureCounts* (Liao *et al.*, 2014) with default settings except that both ends must be aligned and chimeric fragments excluded (i.e. -*B* and -*C* option enabled). Fragments Per Kilobase of transcript per Million mapped reads (FPKM) were used as expression units. Protein-coding region prediction was first performed with *TransDecoder* v5.5.0 (http://transdecoder.github.io) with homology searches against the UniProt90 Reference Clusters (https://www.uniprot.org/) using DIAMOND (Buchfink *et al.*, 2015) and Pfam-A database (http://pfam.xfam.org/) using HMMER (Mistry *et al.*, 2013) to maximize the prediction sensitivity. Additionally, identification of accurate gene sets, removal of assembly redundancies, and protein-coding selection were also performed using *EvidentialGene tr2aacds* pipeline (Don Gilbert, 2013) with a minimum amino acid length of 100 (-MINAA=100). All other settings were default. Transcripts classified as 'okaysets' were retained for further downstream analysis. Functional annotation of the reference transcriptome was performed as follows: 1. Using Mercator4 web application (https://www.plabipd.de/portal/web/guest/mercator4) for assigning mapman BIN categories (Schwacke *et al.*, 2019). 2. Retaining the best Blastx search hit against the plant UniProt sprot database (https://www.uniprot.org/) using DIAMOND with the --*more-sensitive*, --*max-hsps* 1, --*max-target-seqs* 1 option enabled, 3. Incorporation of Pfam domain and homolog annotation from TransDecoder's protein-coding prediction pipeline. Gene space completeness was assessed using Benchmarking Universal Single-Copy Orthologs (BUSCO) v3 (Waterhouse *et al.*, 2018) with default settings except for the lineage database and assessment mode set to -*l* embryophyta_odb10 and -*m* genome, respectively. For *Centranthera*, differential expression between leaf, root, and stem tissues and transcript abundance data (transformed and expressed as FPKM in this study) were obtained from Zhang *et al.* (2019). *Escobedia* orthologs from *Arabidopsis* and *Centranthera* were inferred using CRB-BLAST with default settings (Aubry *et al.*, 2014).

**Methods S3. RT-qPCR.**

mRNA was retrotranscribed into cDNA using oligo-dT as a primer with NZY First-Strand cDNA Synthesis kit (Nzytech). RT-qPCR was performed in a Light Cycler 480 apparatus

(Roche), using SYBRGreen I as a fluorescent reporter and Platinum Taq Polymerase. For 10 µl of quantitative PCR, 2 µl of the cDNA was mixed with 5 µl of 2× SYBRGreen (Roche), 0.3 µm of forward and reverse primers, and 2,4 µl of dH2O. RT-qPCR data were normalized with the actin gene (DN8798_c0_g1_i1). The amplification conditions were as follows: an initial denaturation at 95°C for 10 min, followed by 40 cycles at 95°C for 10 s and 60°C for 30 s. Moreover, melting at 95°C for 2 s, 65°C for 15 sec, and a gradual increment at 95. Three technical replicates for each biological repetition and three biological repetitions were performed.

**Methods S4. Structural and anatomical analyses.**

To determine the localization of orange pigment inside the roots and in the haustoria attached to the host root, fresh samples were incubated in cryoprotectant solution series (sucrose) to cut in a cryostat. Subsequently, samples were embedded in a tissue-freezing medium (Fisher) and frozen up to -19°C. Longitudinal and cross-sections (10-30 µm) were cut in a cryostat Thermo Scientific HM525 NX and then analysed in light and with a Leica TCS SP5 Confocal Laser Scanning Microscope (CLSM). For scanning electron microscopy (SEM) analyses the fixed and dehydrated samples were submitted to critical point drying with $CO_2$ (EM CPD030, Leica). Then, the samples were covered with 30 nm gold using an EMSCD500 high vacuum sputter coater (Leica). Images were taken and recorded using a JEOL XL 30. To analyse the haustorium and root internal structure, fixed and dehydrated samples were infiltrated and embedded with hydroxy ethyl methacrylate (Leica) (Gerrits and Smid, 1983). Sections 5 µm in thickness were cut using a RM 2125RT rotating microtome (Leica). The sections were visualized using a Olympus BX-40 microscope, and images were taken with a Olympus DP71 digital camera. For transmission electron microscopy (TEM), the samples were fixed in 2.5 % glutaraldehyde in 0.1M sodium cacodylate buffer 0.1 M, pH 7.2, and sucrose 0.2 M for 12 h. Furthermore, samples were post-fixed using a solution containing 2% osmium tetroxide and a sodium cacodylate buffer (1:1) for 4 h at room temperature. Finally, samples were washed in a cacodylate buffer, dehydrated in an acetone series, and embedded in Spurr’s resin. Semi-thin 700 nm sections were performed on glass slides to select the region for ultrastructural analysis. Afterward, ultrathin 60 nm sections were contrasted. Images were taken and recorded with a JEOL JEM1011 transmission electron microscope for further analysis.

### Supporting Information - Figures

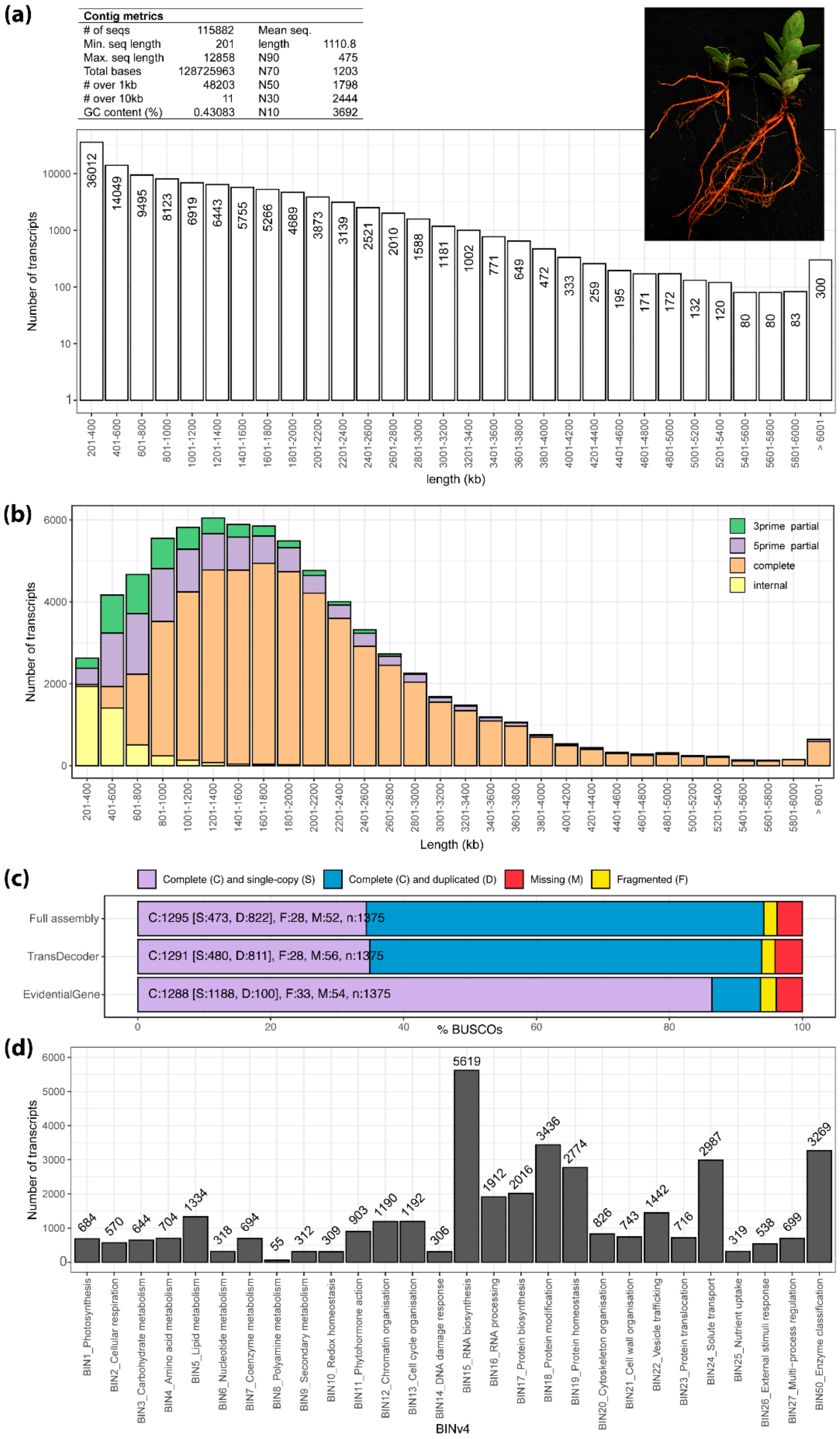

**Fig. S1** *De novo* transcriptome assembly from *Escobedia* roots. **(a)** Summary statistics of the final assembly and length distribution frequency of transcripts (inset shows an image of representative *Escobedia* plants whose roots were used for RNA extraction); **(b)** length distribution frequency of transcripts containing both start and stop codons; **(c)** BUSCO assessments with the embryophyte lineage database; **(d)** functional annotation of the reference transcriptome with MapMan BIN v4 categories.

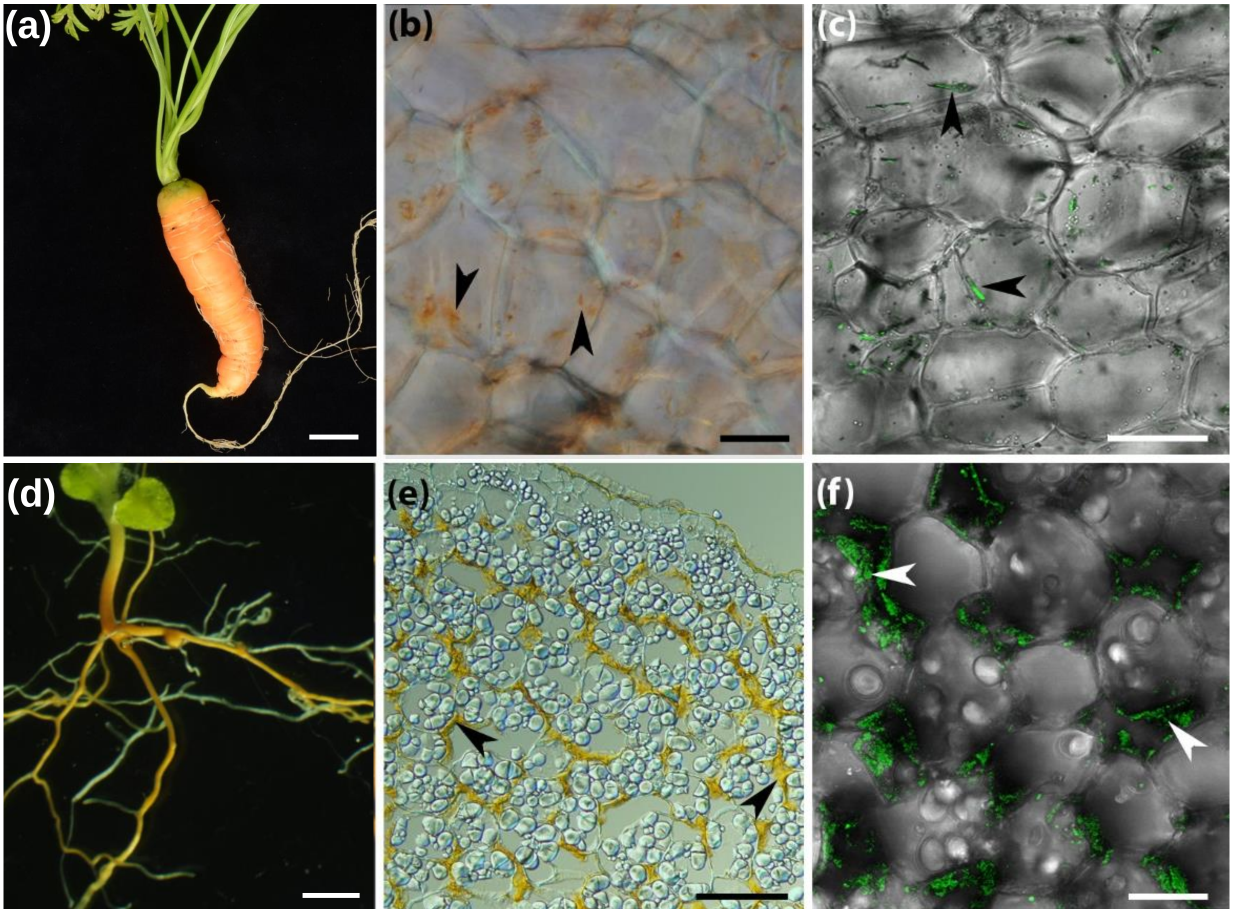


**Fig. S2** Carotenoid and azafrin distribution in carrot and *Escobedia* roots. Panels correspond to samples from carrot **(a-c)** and *Escobedia* **(d-f)**. Roots from carrot **(a)** and *Escobedia* **(d)** plants were collected and used to generate cross-sections for light **(b, e)** and confocal **(c, f)** microscopy. Orange colour in light microscopy sections and green fluorescence in confocal images correspond to carotenoids (in carrot) or azafrin (in *Escobedia*). Arrowheads show chromoplasts of carrot root cells and azafrin in the extracellular space of the *Escobedia* root cortex. Scale bar: (a)=2 cm, (b, c, e)=50 µm, (d)= 2 mm, (f)=20 µm.

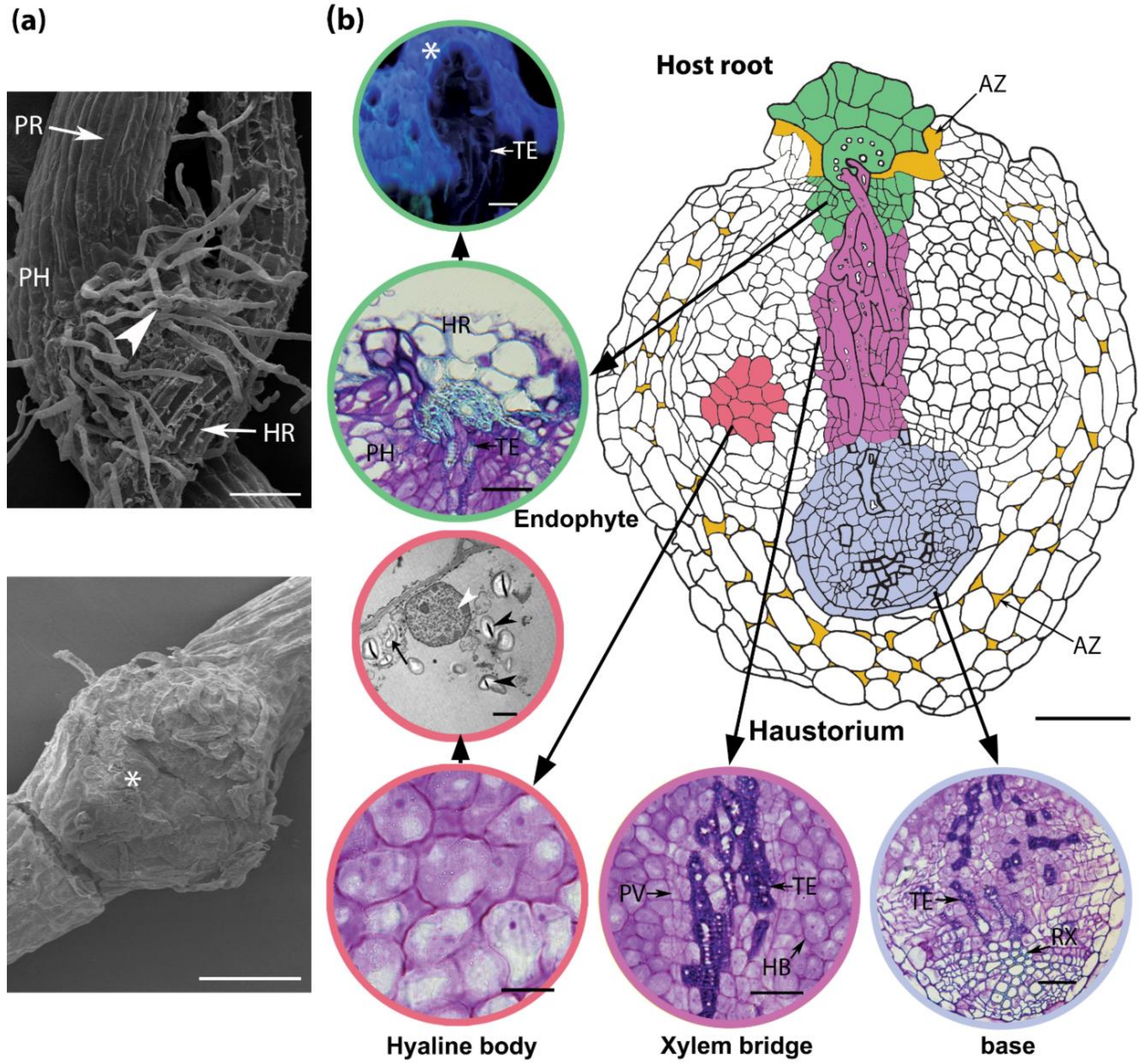


**Fig. S3** Overview of the *Escobedia* haustorium. **(a)** SEM image of haustorium with haustorial hairs (arrowhead) attached to a host root from *Penissetum perpureum* (upper panel) and haustorial opening, which connects with the host root (asterisk, lower panel). Scale bar: 100 μm. **(b)** Illustration showing a longitudinal section of the mature haustorium attaching to the host root. Toluidine blue-stained sections are also shown. The haustorium base (in blue) connects with the host root through the xylem bridge of the haustorium (in purple), consisting of provascular tissue and tracheary elements that grow towards the host root. The hyaline body (in red) shows cells with dense cytoplasm and starch grains (black arrows), Golgi vesicles (narrow arrow), and large nuclei (white arrow). The endophyte is formed by intrusive cells from the haustorium embedded within the host body (in green). Detailed haustorium xylem inside the host's metaxylem (asterisk) indicates parasitism success in the host root. The localization of the azafrin pigment in the apoplast space and in the interface between the haustorium and the host root is depicted in orange. Scale bars are 50 μm except in the illustration (100 μm) and the magnification of hyaline body cells (2 μm). AZ, azafrin; HB, hyaline body; HR, host root; PV, provascular tissue; PH, parasite haustorium; PR, parasite root; RX, root xylem; TE, tracheary elements
